# The P600 and P3 ERP components are linked to the task-evoked pupillary response as a correlate of norepinephrine activity

**DOI:** 10.1101/2023.12.13.571424

**Authors:** Friederike Contier, Isabell Wartenburger, Mathias Weymar, Milena Rabovsky

**Affiliations:** Research Focus Cognitive Sciences, University of Potsdam, Potsdam, Germany; Cognitive Neuroscience Group, Department of Psychology, University of Potsdam, Potsdam, Germany; Patholinguistics / Neurocognition of Language Group, Department of Linguistics, University of Potsdam, Potsdam, Germany; Biological Psychology and Affective Science Group, Department of Psychology, University of Potsdam, Potsdam, Germany

**Keywords:** P600, P300, pupillometry, norepinephrine, LC/NE system

## Abstract

During language comprehension, anomalies and ambiguities in the input typically elicit the P600 event-related potential component. Although traditionally interpreted as a specific signal of combinatorial operations in sentence processing, the component has alternatively been proposed to be a variant of the oddball-sensitive, domain-general P3 component. In particular, both components might reflect phasic norepinephrine release from the locus coeruleus (LC/NE) to motivationally significant stimuli. In this preregistered study, we tested this hypothesis by relating both components to the task-evoked pupillary response, a putative biomarker of LC/NE activity. 36 participants completed a sentence comprehension task (containing 25% morphosyntactic violations) and a non-linguistic oddball task (containing 20% oddballs), while the EEG and pupil size were co-registered. Our results showed that the task-evoked pupillary response and the ERP amplitudes of both components were similarly affected by both experimental tasks. Crucially, the size of the pupillary response – both pupil size and its temporal derivative – predicted the amplitude of both ERP components on a trial-by-trial basis. This pattern of results supports the idea that both, the P3 and the P600, might rely on a shared neural generator and, more specifically, that they may both be linked to phasic NE release. Generally, our findings further stimulate the debate on whether language-related ERPs are indeed specific to linguistic processes or shared across cognitive domains. In the case of the P600, the present results indicate that anomalies during language comprehension might rather initiate transient NE activity in response to rare and motivationally significant stimuli more generally.

## 1 Introduction

Language comprehension often seems effortless, but linguistic input is far from ideal. Listeners are confronted with grammatical errors, false starts, and ambiguities that might impede understanding, increase uncertainty, and necessitate reinterpretation. One important brain response associated with such salient events in the linguistic input is the P600 ERP component. It has been typically observed upon grammaticality violations and ambiguities and has therefore long been considered a purely syntax specific signal of sentence re-analysis or reprocessing (e.g., Friederici, 2002; Hagoort, 2003). However, it is also elicited by semantic implausibility, spelling errors, and violations of narrative structures and even learned, non-linguistic sequences, questioning its syntactic and even language specificity (e.g., Christiansen et al., 2012; Kim & Osterhout, 2005; Münte et al., 1998; Osterhout & Holcomb, 1992; Ryskin et al., 2021; Sitnikova et al., 2008; van de Meerendonk et al., 2009). Thus, the underlying neurocognitive mechanisms of the P600 remain unclear.

Here, we investigated the neurobiological basis of the P600 by linking it to the oddball-sensitive P3 to better understand to what extent both components share a common neural generator, as anomaly detection, saliency, and update of representations are not unique to language processing. Across several other domains, the P3 component, elicited by rare and relevant targets (‘oddballs’) compared to frequent standard stimuli, has been associated with surprise, saliency, and context updating (Donchin, 1981; Nieuwenhuis, Aston-Jones, et al., 2005; Polich, 2007). Parsimoniously, it has thus been suggested that the P600 might be a variant of the P3 (e.g., Coulson et al., 1998; Gunter et al., 1997; Sassenhagen & Fiebach, 2019). P3 latency depends on stimulus complexity (Kutas et al., 1977), which would explain the later onset of the P600 in linguistic contexts. Not only do both components share key properties, such as topography of waveforms and polarity, but both are also most sensitive to stimulus probability and task relevance (see e.g., Leckey & Federmeier, 2019, for an overview). If the two components indeed belong to the same family of late positivities, we could learn more about the P600 from the widely studied P3, but more evidence on their similarities or dissimilarities is needed.

Recently, it has been specifically proposed that both the P3 and P600 reflect phasic norepinephrine (NE) release from the locus coeruleus (LC, Bornkessel-Schlesewsky & Schlesewsky, 2019; Murphy et al., 2011; Nieuwenhuis, Aston-Jones, et al., 2005; Sassenhagen et al., 2014; Sassenhagen & Bornkessel-Schlesewsky, 2015; Vazey et al., 2018). The LC/NE system projects widely throughout the cortex and its phasic responses – transient increases in firing rate – have been associated with unexpected uncertainty, network reset, and neural gain (Dayan & Yu, 2006; Nieuwenhuis, Aston-Jones, et al., 2005; Sara & Bouret, 2012). Conditions for NE phasic responses indeed closely mirror those of the P3 and P600: Responses are related to motivational significance and salience as well as highly sensitive to stimulus probability and task relevance (Nieuwenhuis, Aston-Jones, et al., 2005). Most importantly, there is a strong oddball effect on the activity of the LC/NE system (e.g., Aston-Jones et al., 1997). Further, the latency of these NE responses is tightly locked to the timing of behavioral responses (Bouret & Sara, 2004), which is also the case for the latency of both the P3 (e.g., Kutas et al., 1977; Verleger et al., 2005) and P600 (Sassenhagen et al., 2014; Sassenhagen & Bornkessel-Schlesewsky, 2015). In addition, the LC/NE activity exhibits an “inactive” refractory period after phasic bursts (Usher et al., 1999), which could explain the attentional blink effect on both the P3 (Nieuwenhuis, Holmes, et al., 2005; Woods et al., 1980) and possibly P600 (Batterink & Neville, 2013).

One way to link ERP components to the phasic release of NE is to relate them to the task-evoked pupillary response (TEPR), a potential non-invasive NE marker (Joshi et al., 2016; Reimer et al., 2016; Strauch et al., 2022). Electrophysiological studies with animals and fMRI studies in humans suggest that pupillary responses and LC activity are closely related (e.g., Liu et al., 2017; Murphy et al., 2014; Rajkowski et al., 1993; Sterpenich et al., 2006).

The TEPR is indeed also evoked under similar conditions as the late positive ERP components: Importantly, its size is highly dependent on stimulus probability (e.g., Friedman et al., 1973; Qiyuan et al., 1985; Wetzel et al., 2016). Further, previous studies co-registering EEG and the TEPR found an oddball effect on both measures within the same paradigm, although they had difficulties uncovering a direct relationship between the two (Hong et al., 2014; Kamp & Donchin, 2015; LoTemplio et al., 2020; Murphy et al., 2011). One reason why it might be difficult to detect such a correlation in oddball paradigms might be that the TEPR, especially measured as pupil size across the trial, might not only be influenced by the cognitive ‘oddball’ processing which is also reflected in the P3, but additionally influenced by, for instance, later processes of response preparation. It is thus worth further investigating their link, and specifically their exact temporal relationship more closely. Lastly, in language processing, just like the P600, the TEPR is also sensitive to syntactic ambiguity (Engelhardt et al., 2010), complexity (Just & Carpenter, 1993; Schluroff, 1982), and violations (Aydın & Uzun, 2022), but the direct relationship between the P600 and the TEPR has not been studied.

We here systematically investigated the relationship between the late positivities – P3 and P600 – to the LC/NE-system by relating them to the trial-by-trial task-evoked pupillary response. Participants performed both an active sentence processing task in which we manipulated grammaticality (to elicit the P600) and an active oddball task (to elicit the P3) while we co-registered EEG and pupil size. If not only the TEPR, but also the P3 and P600 reflect phasic NE responses, both components’ amplitudes and the TEPR should be larger on syntactic violations and oddballs compared to correct control sentences and standards.

Further, the two measures should be positively correlated on a trial-by-trial basis in both tasks. Just like previous studies, our primary measure for this TEPR was the size of the pupil within a time window post-stimulus onset. In contrast to previous studies, we additionally explored these effects using the temporal derivative of the pupil (McGinley et al., 2015; Murphy et al., 2021; Reimer et al., 2014, 2016; Yang et al., 2020). Since pupil size itself is known to change slowly, the additional advantage of this approach is that it is sensitive to changes on a faster timescale and that it can differentiate between periods of actual dilation and constriction within one response. Throughout the paper, we will use the acronym TEPR to refer to the pupillary response in general, and “pupil size” or “pupil derivative” to refer to either measure specifically. The supplementary material contains analyses of the relationship between ERPs as well as TEPR to the baseline size of the pupil (as a potential marker of tonic NE) in both tasks. Here, we focus on the potential link between the two task-evoked signals (ERP component and TEPR) during oddball and sentence processing.

## 2 Method

### 2.1 Participants

The study was preregistered on the Open Science Framework (OSF, https://osf.io/sm78w/?view_only=2aba21018b994d249a64483953b06021) and potential deviations are stated as such^1^. Analyses code and data are also available on OSF (https://osf.io/xmszg/?view_only=46d4f529d81340fb90c872f5a342bc53). We tested 45 German native speakers with normal or corrected vision (lenses) and no reported history of neurological or psychiatric disorder or language impairments. All participants were right-handed as assessed via the Edinburgh Handedness Inventory (Oldfield, 1971; 12-item version). They received monetary compensation or course credit. From the collected sample, datasets from five participants had to be excluded due to technical errors. Data from four additional participants had to be excluded since they did not contribute enough data points after artifact rejection (at least half of the trials in total and per condition for each task in both measures combined, see preregistration). As preregistered, datasets from 36 participants (30 female, 6 male; mean age: 23.33 years, range: 19-32; mean laterality quotient: 81.6, range: 50-100) remained for analyses. Each individual provided written informed consent for a protocol approved by the ethics committee of the host institution.

### 2.2 Procedure & materials

The experiment took place in an electrically shielded room with ambient light (two battery-powered desk lamps turned away from the participant, appr. 100 lumen). All participants took part in both a visual sentence processing as well as an oddball task. The order of the two tasks was counterbalanced across participants. Stimuli were presented visually on an LED monitor using MATLAB’s PsychToolBox version 3.0 (Brainard, 1997). Participants were positioned within 60cm distance of the screen. All stimuli were presented with low contrast (dark grey font on medium grey background). To additionally reduce the light reflex of the pupil to stimulus on- and offsets during rapid serial visual presentation, experimental stimuli (single letters in oddball task and single words in sentence processing task) were enclosed by a hashtag mask. During inter-stimulus intervals (ISI), the mask was shown, which consisted entirely of hashtags with the same number of characters. Since characters were presented in monospaced font (Courier), this resulted in a continuous, same-sized mask throughout the entire task (of three characters in the oddball and 17 characters in the sentence processing task, see Figure 1).

**Figure 1.**
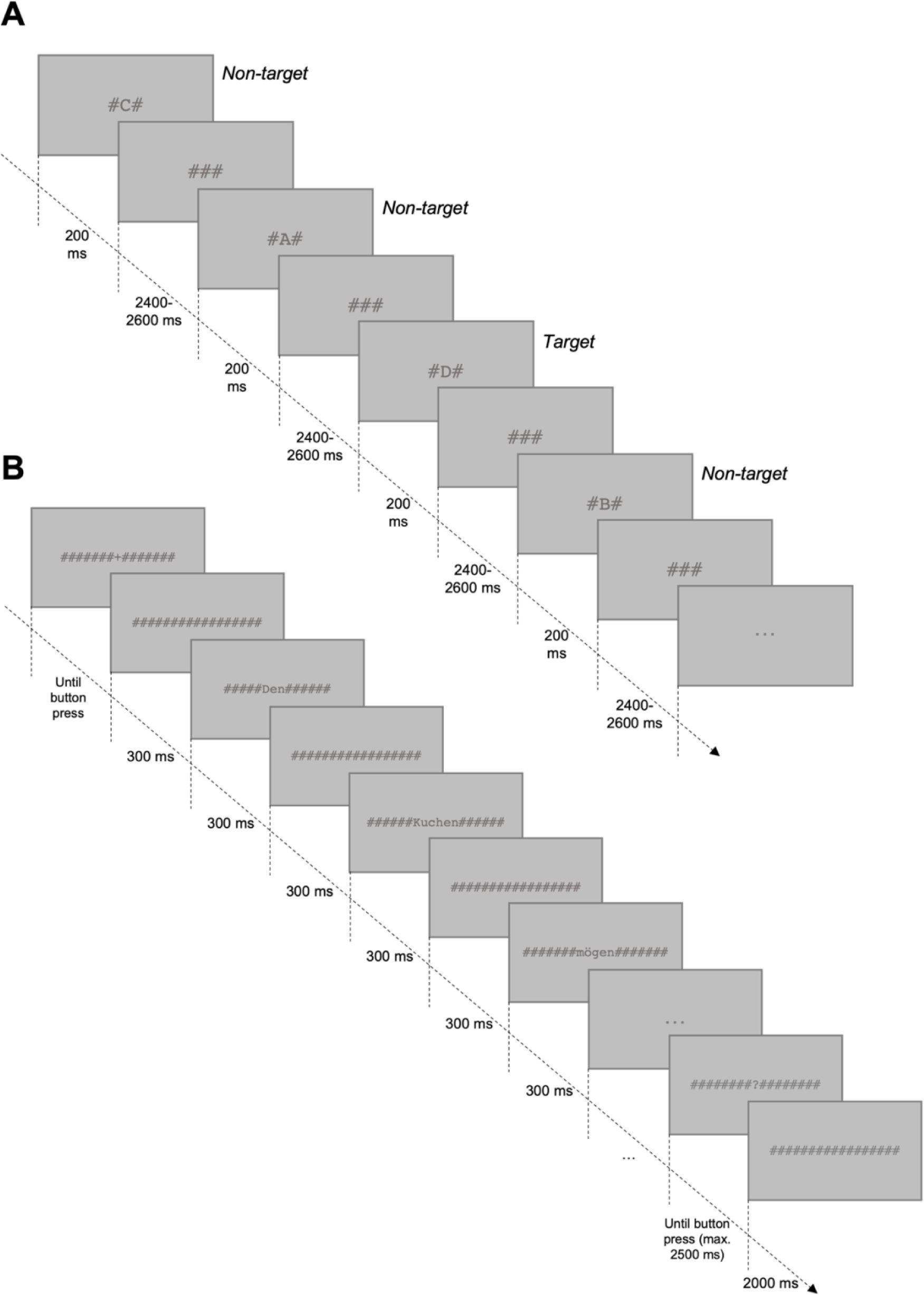
**A**: Stimulus presentation in the oddball task, with an example stimulus sequence in the block in which “D” was the target letter. **B**: Word-by-word stimulus presentation of one trial in the sentence processing task.

#### 2.2.1 Oddball task

We used a modified version of the active visual oddball task by Kappenman et al. (2021), which was designed to elicit a robust P3 component and to reduce as many sensory confounds as possible. The participant’s task is to identify rare target stimuli (“oddballs”) among frequent non-targets (“standards”).

In the task, the letters A-E served as stimuli, which were each presented with a probability of .2. The task comprised five blocks of trials and for each block, a different letter was designated to be the target for that block. Thus, the probability of the target was always .2 but the same physical stimulus served as a target in some blocks and as a non-target in others. Within a block, letters were presented in random order, with the restriction that targets were never followed by targets. The order of blocks was also randomized across participants. We presented a total of 400 trials, divided into five blocks of 80 trials each.

Stimulus letters were presented at the center of the screen (see Figure 1A). As noted above, letters were enclosed by two hashtags, together yielding a size of 2 x 3.8 degrees of visual angle. Stimulus letters were presented for 200 ms. Since the pupil response takes some time to return to baseline, we used a long ISI of 2400-2600 ms (showing a mask of three hashtags). ISIs were jittered in steps of the screen refresh interval (16.67 ms) using a rectangular distribution.

Participants were instructed to respond as quickly and accurately as possible whether each letter was the target or a non-target in each block. There was no designated time window for responses. Participants could respond any time between onset of the stimulus letter and the end of the long ISI. Two marked (ctrl) keys on the computer keyboard served as response buttons, each pressed by the index finger of the respective hand. Stimulus-response mapping was counterbalanced across participants and held constant for both tasks within a participant (i.e., the same key, either the left or right ctrl key, for targets in the oddball task and grammatical violations in sentence processing task and vice versa).

Written instructions were presented at the beginning of the task and reminder instructions, indicating the target letter in the current block, were presented after each break. Experimental trials were preceded by 27 practice trials with a different set of letters than the experimental stimuli. Participant-controlled rest breaks were provided between blocks and after half of the trials within each block. The task lasted approximately 25 minutes.

#### 2.2.2 Sentence processing task

We used a classic sentence processing task in which sentences are presented word by word and participants are required to make a grammaticality judgement after each sentence. 75% of the sentences were correct sentences and ¼ contained morphosyntactic violations, which have been shown to robustly elicit a P600 response (Osterhout & Mobley, 1995).

German sentences by Sassenhagen & Fiebach (2019, Exp. 2) served as the basis for our stimulus set. These sentences were simple, plausible subject-verb-object (S-V-O) or object-verb-subject (O-V-S) sentences. A second version of each sentence with an agreement violation had been constructed by exchanging the verb or pronominal subject for a morphosyntactically ill-fitting one from another sentence. In the original set, target word class and position systematically differed between sentence types: In S-V-O sentences, the target word was a verb in the 3^rd^ position (e.g., “Der Garten <u>blüht</u> schon im Frühling / Die Gärten *<u>blüht</u> schon im Frühling.” [The[sg] garden[sg] <u>blossoms</u>[sg] already in spring. / The[pl] gardens[pl] *<u>blossoms</u>[sg] already in spring.]). In O-V-S sentences, the target word was a pronominal subject in the 4^th^ position (Den Kuchen mögen <u>sie/*er</u> ganz besonders. [The[sg] cake[sg] like[pl] <u>them</u>[pl]<u>/*he</u>[sg] very much, lit. They/*he particularly like the cake.]). Pupil size might differ between positions in the sentence, generally increasing or decreasing over the course of a sentence. To avoid that pupil size might then also systematically differ between sentence types and target word class, and hence pose confounds in our results, we aligned the target word position to the 4^th^ position by adding a modifier before the subject noun in all S-V-O sentences (e.g., “Der neue Kellner <u>steht</u> hinter dem Tresen. / Die neuen Kellner *<u>steht</u> hinter dem Tresen.” [The[sg] new[sg] waiter[sg] <u>stands</u>[sg] behind the counter. / The[pl] new[pl] waiters[pl] *<u>stands</u>[sg] behind the counter.]). We also added more sentences in the same fashion, resulting in a final set of 240 sentences, of which 60 served as correct fillers (added to increase the proportion of correct sentences) and 180 as experimental sentences. We created three lists, each including the 60 fillers, 60 of the experimental sentences with a morphosyntactic violation (violation condition) and 120 correct experimental sentences (i.e., sentences without a morphosyntactic violations; henceforth: control condition). Thus, each experimental sentence appeared in one list as a violation and in the other two as a correct control. Including the correct fillers (which were not part of the analysis), each participant was presented with 240 sentences of which 25% contained a violation.

Sentences were presented in random order, with the only constraint that violation sentences were never directly followed by another violation sentence. Each trial (see Figure 1B) started with a fixation cross, which was shown until the participant pressed the space bar. Words were then presented one after another, each for 300 ms with a 300 ms inter-word interval. If a non-target word’s length exceeded nine letters, 33.33 ms were added to its presentation duration for each additional letter (for comparable approaches, see e.g., Hodapp & Rabovsky, 2021; Nicenboim et al., 2020). 600 ms after the offset of the final word, a question mark appeared, prompting the participant to decide whether the presented sentence was grammatically correct or incorrect. Two marked (ctrl) keys on the computer keyboard served as response buttons and stimulus-response mapping was counterbalanced across participants but held constant for both tasks within each participant (see above). The question mark was shown until button press or 2500 ms had passed. The inter-trial interval lasted 2000 ms.

Words were enclosed in a hashtag mask (with at least one hashtag in the beginning and one in the end) with the number of hashtags adjusted to word length. This resulted in a same sized mask of 17 characters across all words (see Figure 2B). The prompts (fixation cross and question mark) were also enclosed by the hashtag mask (17 characters in total). Likewise, in between words and prompts, 17 hashtags were shown, so that participants were presented with a continuous, same sized mask throughout the entire task, minimizing luminance contrasts at stimulus on- and offsets. The mask had a size of 1.4 x 17.1 degrees of visual angle.

**Figure 2.**
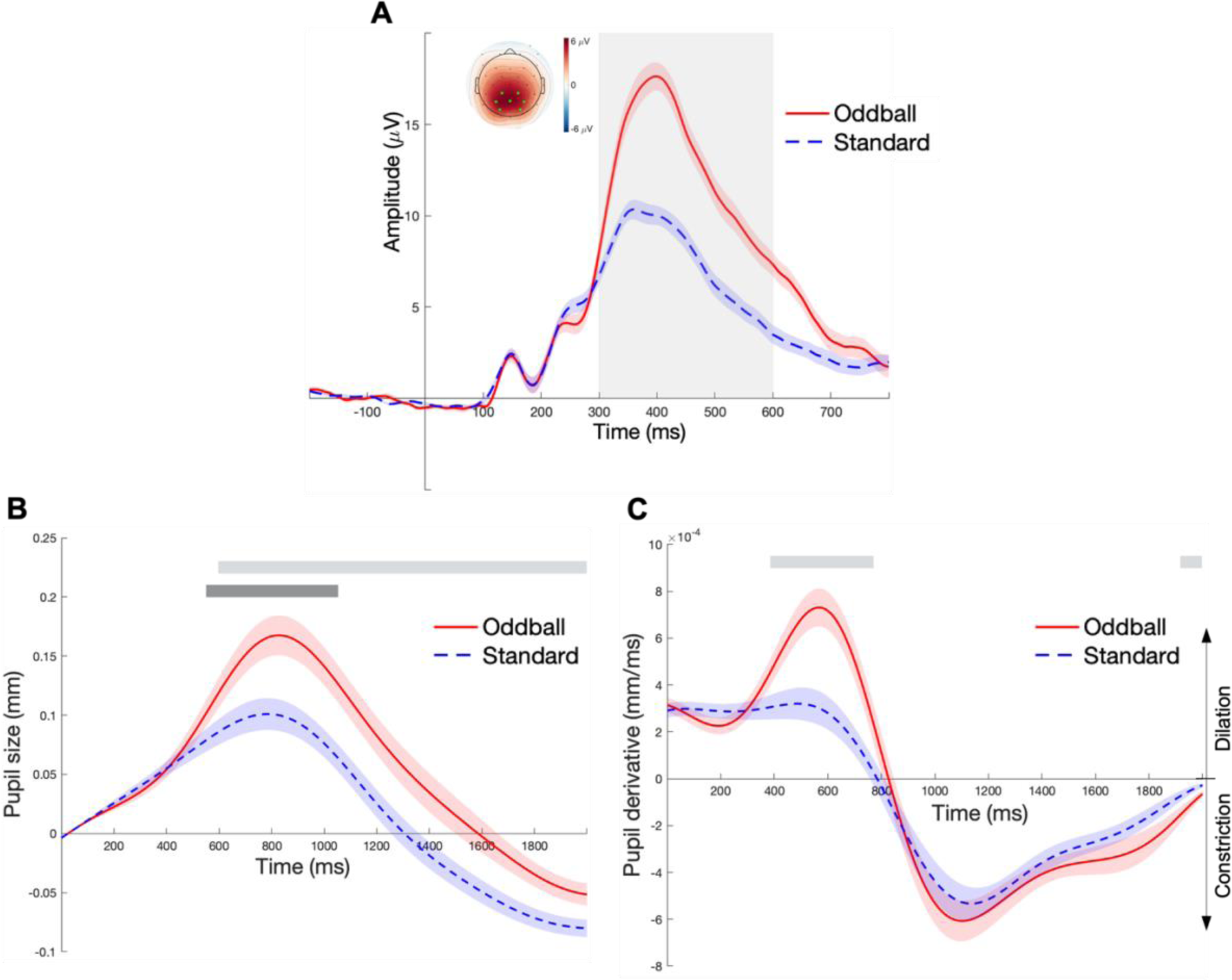
Grand averaged ERP amplitude (**A**), pupil size (**B**), and pupil derivative (**C**) for the two conditions in the oddball task. **A**: P3 amplitudes were calculated as the mean in the time interval marked by the grey box (300-600 ms) within the posterior channel cluster marked in light green in the topographic map. The topographic map shows the mean amplitude difference between oddball and standard stimuli within the same time window. **B**: The light grey bar highlights the time cluster for which there was a significant difference in pupil size between the two conditions as determined using the cluster-based permutation test (598-2000 ms). This time window was used to compute mean pupil size values for each trial in our initial analysis. The dark grey bar highlights the time window around the peak of the dilation (552 - 1052 ms), which was used to compute mean pupil size values for each trial for the exploratory model whose results are depicted in Fig. 3A. **C**: The light grey bar highlights the time cluster for which there was a significant difference in pupil derivative between the two conditions as determined using the cluster-based permutation test (dilation at 386-770 ms and constriction at 1918-2000 ms). The lefthand time window (dilation) was used to compute mean pupil derivative (i.e., dilation) values for each trial in our statistical models. Error bands in A, B, and C indicate the standard error of the group mean (SEM) based on subject-wise average time series.

Participant-controlled rest breaks were provided between every 30 trials. Before the start of the experimental sentences, participants were given written instructions and practiced the task with eight additional sentences. The task lasted around 40 minutes.

### 2.3 Data acquisition & processing

#### 2.3.1 Pupil data

Pupil size from both eyes was recorded at a Tobii Pro X3-120 eye tracker with a sampling rate of 40 Hz. Before each task, participants completed a classic 5-point calibration procedure (see e.g., Mckinnon et al., 2020; Trueswell et al., 2013). Because the raw data contained high frequency jitter, they were first smoothed using a Savitzky-Golay filter (window size = 20 datapoints, polynomial degree of 2). Epochs (0-2000 ms relative to stimulus/target word onset) were preprocessed using the pipeline by Kret & Sjak-Shie (2019), which removes dilation speed outliers, trend-line deviations, and secluded samples (e.g., due to noise of erroneous pupil detection during shut eyes). Data from the left and right pupil were then averaged. At this point, epochs were excluded in which 50% or more datapoints were missing. Remaining trial data were upsampled to the same rate as the EEG data (500 Hz) and low-pass filtered (zero-phase, cutoff at 4 Hz). We then additionally excluded epochs with missing consecutive datapoints (gaps) of > 50% of datapoints within an approximated temporal region of interest for the task-evoked dilation (oddball: 600-1200 ms; sentence processing task: 900-1500 ms). Missing values in epochs that had survived this threshold were interpolated using a cubic spline interpolation and data were baseline corrected to an interval 0-50 ms relative to stimulus/target word onset (Mathôt & Vilotijević, 2022). Pupil derivative was calculated for each timepoint within an epoch by using MATLAB’s ‘diff’ function (Mäki-Marttunen, 2021; van den Brink et al., 2016). For each trial, this resulted in a derivative time series indicating how fast the pupil changes its size at each time point, with positive values indicating dilation and negative values indicating constriction (see Fig. 2C and 4C). Finally, pupil trial data in the following cases were excluded: 1) with inaccurate responses, 2) with responses earlier than 150 ms after stimulus onset in the oddball task, 3) with excluded EEG data due to artifacts, and 4) control trials following violation trials (sentence processing task) and standard trials following oddball trials (oddball task).

As there is no established temporal region of interest for the dilation within the TEPR in either oddball or sentence processing task, we preregistered to define this time window as the significant cluster of condition differences as indicated by a nonparametric cluster-based permutation test (Maris & Oostenveld, 2007). Note that such tests only reliably identify significant cluster sizes but not single time points included in the cluster, such as the exact start and end point of condition differences (Sassenhagen & Draschkow, 2019). However, we still used these time windows as a close temporal approximation to potential time windows in which the pupil size in each task differed between conditions and possibly related to the ERP amplitude. We used the clusterPerm package in R (Barr, 2020) and performed tests on aggregated data with 1000 permutations. The cluster-based permutation test generally indicated a difference of pupil size between conditions in both tasks (both *p* < .001). The corresponding clusters in the observed data were as follows: 598-2000 ms in the oddball task (light grey bar in Fig. 2B) and 730-1622 ms in the sentence processing task (light grey bar in Fig. 4B). Thus, for each trial, pupil size as a measure in the statistical analyses below was quantified as the mean pupil size in these time windows. The derivative of the pupil on the other hand revealed that – in contrast to standards and controls – oddballs and violations first dilated, then constricted, resulting in two time clusters with significant differences between conditions in each task (light grey bars in Fig 2C and 4C). In the oddball task, the oddball dilation was at 386-770 ms (*p* < .001) and the late constriction at 1918-2000 ms (*p* = .007). In the sentence processing task, violations dilated more than controls at 530-1078 ms (*p* < .001) and then constricted more than controls from 1170-2000 ms (*p* < .001). Since we were primarily interested in the relationship between ERP amplitudes and the dilatory response of the pupil, our trial-wise pupil derivative values for subsequent analyses are based on the mean derivative within the time window of the respective dilation (i.e., the first time window in both tasks).

#### 2.3.2 EEG data

Participants’ electroencephalogram (EEG) was recorded using 32 active Ag/AgCl electrodes (actiCHamp system, Brain Products GmbH, Gilching, Germany), spaced according to the international 10-20 system. Impedances were reduced below 10 kOhm if possible. The ground electrode was positioned at the forehead and the reference at the left mastoid. Bilateral VEOG as well as HEOG electrodes (right side) were placed to capture eye movements and blinks. EEG was recorded at a sampling rate of 1000 Hz (bandpass filter of .016-250 Hz, time constant of 10s).

Offline, data were preprocessed in Matlab R2020a, using the EEGLAB (Delorme & Makeig, 2004) toolbox. Data were downsampled to 500 Hz and re-referenced to the average of the left and right mastoid. Ocular artifacts were corrected using independent component analysis (Infomax ICA, Jung et al., 2001). ICA was trained on segments spanning -1 to +2s (sentence processing task) and -0.5 to +2s (oddball task) post stimulus onset, after artifact rejection (removal of epochs exceeding +150/-150 mV), which were extracted from continuous data filtered with a 1-30 Hz Butterworth filter. Independent components were removed via the IClabel plug-in (Pion-Tonachini et al., 2019), which identified eye- contaminated segments with a probability > 30%. We additionally removed (max. three) channel noise components based on visual inspection.

Corrected, continuous data were high-pass filtered (0.1 Hz, two-pass Butterworth with a 12 dB/oct roll-off) and low-pass filtered (30 Hz, two-pass Butterworth with a 24 dB/oct roll-off). Line noise caused by the set-up being close to the power supply for the PC and eye-tracker was filtered out using EEGLAB’s CleanLine plug-in (at 50/100 Hz). Bad channels (exceeding an absolute z-score of 3 regarding voltage across channels) were interpolated using a spherical spline function. We then epoched the data from -200 to 1100 ms (sentence processing task) and from -200 to 800 ms (oddball task), time-locked to target word/stimulus onset. Epoched data were baseline corrected relative to the 200 ms interval preceding the onset of the stimulus/target word. We automatically removed artifactual trials exceeding +/-75 mV.

We additionally excluded epochs related to incorrect or too early responses (150 ms, oddball task), standards following oddballs (oddball task) and control trials after violation trials (sentence processing task), or because of artifactual or missing pupil data. On average, participants provided 53 violation sentence trials (range: 38-60), 73 control sentence trials (range: 61-82), 67 oddball trials (range: 43-77) and 208 standard trials (range: 153-234). Data for both components were analyzed for a parietal region of interest (ROI: CP1, CP2, P3, Pz, P4, PO3, PO4) within a 300-600 ms (P3) and 600-900 ms (P600) time window. Spatial ROIs and time windows most closely resembled the topography and time where each of the components had been shown to be largest in the respective task (Kappenman et al., 2021; Sassenhagen & Fiebach, 2019).

### 2.4 Statistical analyses

We performed linear mixed-effects model (LMM) analyses using the package lme4 as implemented in *R* (R core Team, 2018). All models were built using trial-by-trial data. We first tested for the effect of the respective manipulation on different measures in the oddball task (1) and sentence processing task (2):

1. *Accuracy/RT/P3/TEPR ∼ condition + (1 + Accuracy/RT/P3/TEPR | participant)*
2. *Accuracy/RT/P600/TEPR ∼ condition + (1 + Accuracy/RT/P600/TEPR | participant) + (1 + Accuracy/RT/P3/TEPR | sentence)*

In these models, condition was sum coded (Schad et al., 2018; Oddball task: Oddball: - 0.5, standard: 0.5,; Sentence processing task: violation: -0.5, control sentence: 0.5). The central models testing the relationship between the two measures assumed to reflect NE activity had the following structure in the in the oddball task (3) and sentence processing task (4):

1. *P3 ∼ TEPR + (1 + TEPR | participant)*
2. *P600 ∼ TEPR + (1 + TEPR | participant) + (1 + TEPR | sentence)*

These models were run with only data from the respective target condition (oddball/violation) and, as additionally preregistered, with data from both respective conditions together to increase power and variance. As additional, exploratory analyses, we also ran a version of the main ERP-pupil size models including condition as a covariate to control for the shared oddball/violation effect. However, assuming that the TEPR and ERP component are both reflecting a latent variable (NE) that is most sensitive to the oddball/violation effect, the relationship between the two measures is inherently tied to variance caused by the oddball/violation effect (see the discussion for more on this reasoning), rendering these control models less reliable.

All models included a random effects structure (see equations above). Following the recommendations by Barr et al. (2013), we tried to fit this maximal random effect structure but reduced its complexity successively until the model converged. As pre-registered, we first increased the number of iterations to 10000, then introduced an optimizer (“bobyca”), and then removed correlations between random intercepts and slopes and if necessary, entire random slopes. Almost all models could retain random slopes by subjects for the predictor of interest, those that could not, are marked as “RSI model” (random subject intercept model).

To help model convergence, TEPR and ERP values were z-scored (within participant and task). RTs were log-transformed, as they were extremely positively skewed (oddball: 2.09, sentence processing: 2.331). The significance of fixed effects was determined via likelihood ratio tests to compare the fit of the model to that of a reduced model lacking the respective fixed effect but including all remaining fixed effects as well as the same random effect structure. As pre-registered, we used the standard *p* < .05 criterion for determining if the likelihood ratio tests suggest that the results are significantly different from those expected if the null hypothesis were correct.

## 3 Results

### 3.1 Oddball task

#### 3.1.1 Behavioral performance

Overall accuracy was very high (*M* = 97%; *SD* = 3%) and was slightly better in the standard condition (*M* = 98%, *SD* = 2%) than in the oddball condition (*M* = 91%, *SD* = 7%; *β* = -2.505, *SE* = 0.207, *z* = -12.13, *p* < .001). Within correct trials, responses to frequent stimuli were faster than to oddball stimuli, replicating previous findings (e.g., Strauch et al., 2020).

This pattern was found both within behavioral responses overall (*β* = 0.081, *SE* = 0.011, *t* = 7.507, *χ2* = 34.63, *p* < .001) as well as within the subset of trials used for the subsequent P3 and TEPR analyses (see Methods section for trial exclusion criteria; *β* = 0.072, *SE* = 0.011, *t* = 6.615, *χ2* = 29.311, *p* < .001). Note that all behavioral results in both tasks are based on additional, exploratory analyses (i.e., not pre-registered).

#### 3.1.2 Effect of condition on ERP amplitude and TEPR

Condition means of the ERP amplitude, pupil size, and pupil derivative are shown in Figure 2A, 2B, and 2C, respectively. Replicating the classic P3 effect, we found that ERP amplitudes over posterior regions were significantly larger for oddballs than standards (*β* = 0.270, *SE* = 0.017, *t* = 15.46, *χ2* = 74.29, *p* < .001). Similarly, within the time windows indicated by the cluster-based permutation tests, the pupillary response was significantly larger in response to oddballs than standards, as there was a significant condition effect on both pupil size (*β* = 0.178, *SE* = 0.019, *t* = 9.264, *χ2* = 43.707, *p* < .001) and pupil derivative (*β* = 0.001, *SE* = 0.0001, *t* = 13.480, *χ2* = 65.443, *p* < .001, additional, exploratory analysis).

Thus, the oddball manipulation affected the ERP amplitude and the TEPR in the same way, increasing upon rare and behaviorally relevant oddballs compared to frequent standard stimuli.

#### 3.1.3 Relationship between ERP amplitude and TEPR

To investigate whether the P3 and pupillary response might be linked to a common neural generator, we analyzed whether ERP amplitudes were positively predicted by the pupillary response on a trial-by-trial basis. We pre-registered to analyze the relationship between the P3 time window for ERPs (300-600ms) and the time window revealed by the cluster-based permutation test for the pupil size (598-2000 ms). However, this latter time window regarding the pupil size was surprisingly long, spanning over almost three quarters of the epoch (598-2000 ms). Consequently, the main analyses failed to find any relationship between the P3 and pupil size averaged within this long time window (within oddballs: *β* = - 0.017, *SE* = 0.012, *t* = -1.407, *χ2* = 1.915, *p* = .166, see Fig. 3A; across both conditions: *β* = 0.012, *SE* = 0.009, *t* = 1.416, *χ2* = 2.023, *p* = .155). To better capture the main part of the response, we therefore conducted an additional exploratory analysis in which we restricted the pupil size time window to a 500 ms window centered around the peak of the pupil response at 802 ms. In this shorter window, there was a statistically significant, positive relationship between the two measures of interest across both conditions (*β* = 0.068, *SE* = 0.010, *t* = 6.61, *χ2* = 31.427, *p* < .001; see Fig. 3B; but not within only oddballs: *β* = 0.013, *SE* = 0.016, *t* = 0.791, *χ2* = 0.621, *p* = .431). Crucially, in the analysis including both conditions, this relationship persisted even when controlling for the effect of condition (*β* = 0.018, *SE* = 0.007, *t* = 2.367, *χ2* = 5.357, *p* = .021; additional exploratory model). In the supplementary material, we provide additional, preregistered analyses decomposing the pupillary response using PCA that further support the focus on this peak of the pupil response. In particular, we demonstrate that trial-by-trial P3 amplitude was exclusively predicted by an early part of the pupillary response, that is, a principal component that already peaked around 700-800 ms after stimulus onset, in analyses focusing on oddball trials only.

**Figure 3.**
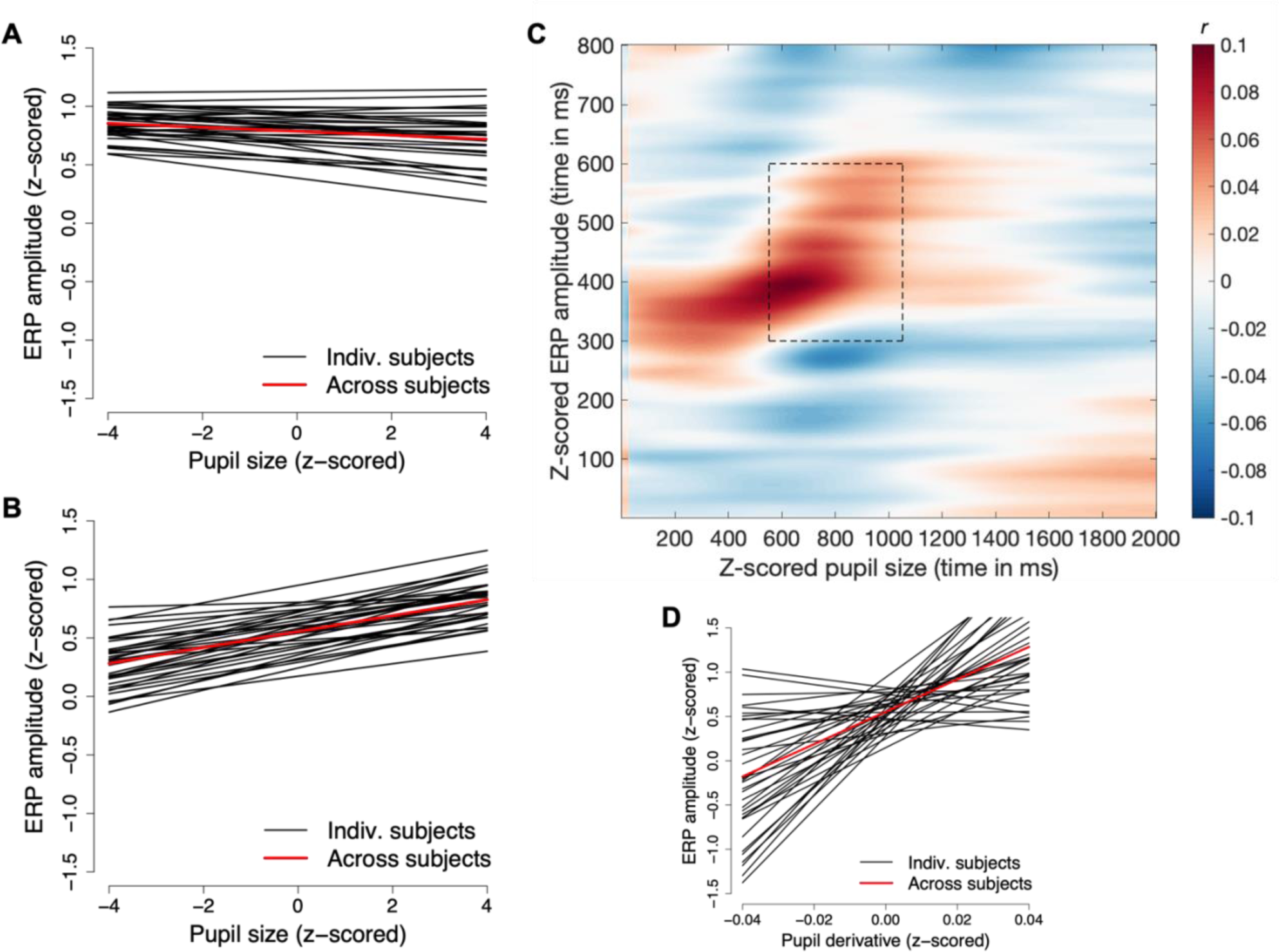
Relationship between the P3 and the TEPR in the oddball task. **A**: Predicted values for each participant and their mean from the main preregistered model testing the relationship between P3 amplitude and mean pupil size within the long time window determined via the CBPT (light grey bar in Fig. 2B) within trials of the oddball condition. **B**: Predicted values for each participant and their mean from the additional, exploratory model testing the relationship between P3 amplitude and mean pupil size around the peak of the dilation (dark grey bar in Fig. 2B) within trials of both conditions. **C**: Cross-correlation of P3 amplitude and pupil size over time (average R of correlations within participants). The dashed rectangle marks the combined time window which was used for the model in B. **D**: Predicted values for each participant and their mean from the model testing the relationship between P3 amplitude and mean pupil derivative within trials of both conditions (in dilation time window indicated by the lefthand light grey bar in Fig. 2C).

We additionally provide a visualization of how ERP amplitudes at each time point correlate with pupil size at each time point over the entire epoch across both conditions within each participant (Fig. 3C displays the mean correlation across participants). Note that this cross-correlation is intended to visually demonstrate the temporal specificity of the relationship between the two measures, while we refrain from additional inferential statistics here to avoid multiple testing. This cross-correlation plot confirms that 1) the correlation between the two measures is indeed strongest in an earlier time window, around the peak of the pupillary response, and 2) that the later part of the pupillary response is not or even negatively correlated with P3 amplitude, thus diminishing any effect when taking the pupil mean across a long time window. Figure A2 in the supplementary material shows these cross-correlations over time for trials from each condition separately. They additionally demonstrate that trials from the oddball, not standard condition are driving this early, positive correlation between ERP amplitude and pupil size.

Crucially, in contrast to pupil size, pupil derivative analyses showed a much earlier and shorter time period in which the pupil dilated more to oddball than standard stimuli (light grey cluster bar in Fig. 2C, add. exploratory analyses). Within this time window derived from the cluster-based permutation test, pupil derivative also displayed a positive relationship to P3 amplitude on a trial-by-trial basis considering trials from both conditions (*β* = 18.3, *SE* = 3.5433, *t* = 5.165, *χ2* = 21.619, *p* < .001; see Fig. 3D; but not within oddballs: *β* = 6.641, *SE* = 4.38, *t* = 1.513, *χ2* = 2.307, *p* = .129). Taken together, this suggests an early, temporally specific correlation between the pupillary response and the P3 within an oddball task

### 3.2 Sentence processing task

#### 3.2.1 Behavioral performance

As in the oddball task, accuracy in the grammaticality judgment was very high (*M* = 96%; *SD* = 4%), but did not differ between violations and controls (*β* = -0.163, *SE* = 0.227, *z* = -0.72, *χ2* = 47.979, *p* = .472), indicating that participants attentively read both types of sentences. In correctly identified sentences, judgements were faster following violation than control sentences. This was the case in judgements overall (*β* = -0.243, *SE* = 0.035, *t* = -6.905, *χ2* = 47.501, *p* < .001, RSI model) as well as within the subset of trials used for the subsequent P600 and TEPR analyses (see Methods section for trial exclusion criteria; *β* = - 0.251, *SE* = 0.062, *t* = -4.044, *χ2* = 13.82, *p* < .001).

#### 3.2.2 Effect of condition on ERP amplitudes and TEPR

Condition means of the ERP amplitude, pupil size, and pupil derivative are shown in Figure 4A, 4B, and 4C, respectively. Violations generally elicited a late, positive ERP amplitude deflection over posterior electrodes and a large dilation of the pupil. As expected, we found that ERP amplitudes were significantly larger in response to targets in violation than control sentences (*β* = 0.372, *SE* = 0.029, *t* = 12.988, *χ2* = 63.678, *p* < .001), reflecting the classic P600 effect. Similarly, the pupil dilated significantly more in response to violations than controls, which was evident in both pupil size (*β* = 0.207, *SE* = 0.034, *t* = 6.165, *χ2* = 26.333, *p* < .001) and pupil derivative (*β* = 0.001, *SE* = 0.0001, *t* = 8.771, *χ2* = 42.16, *p* < .001, add. exploratory analysis).

**Figure 4.**
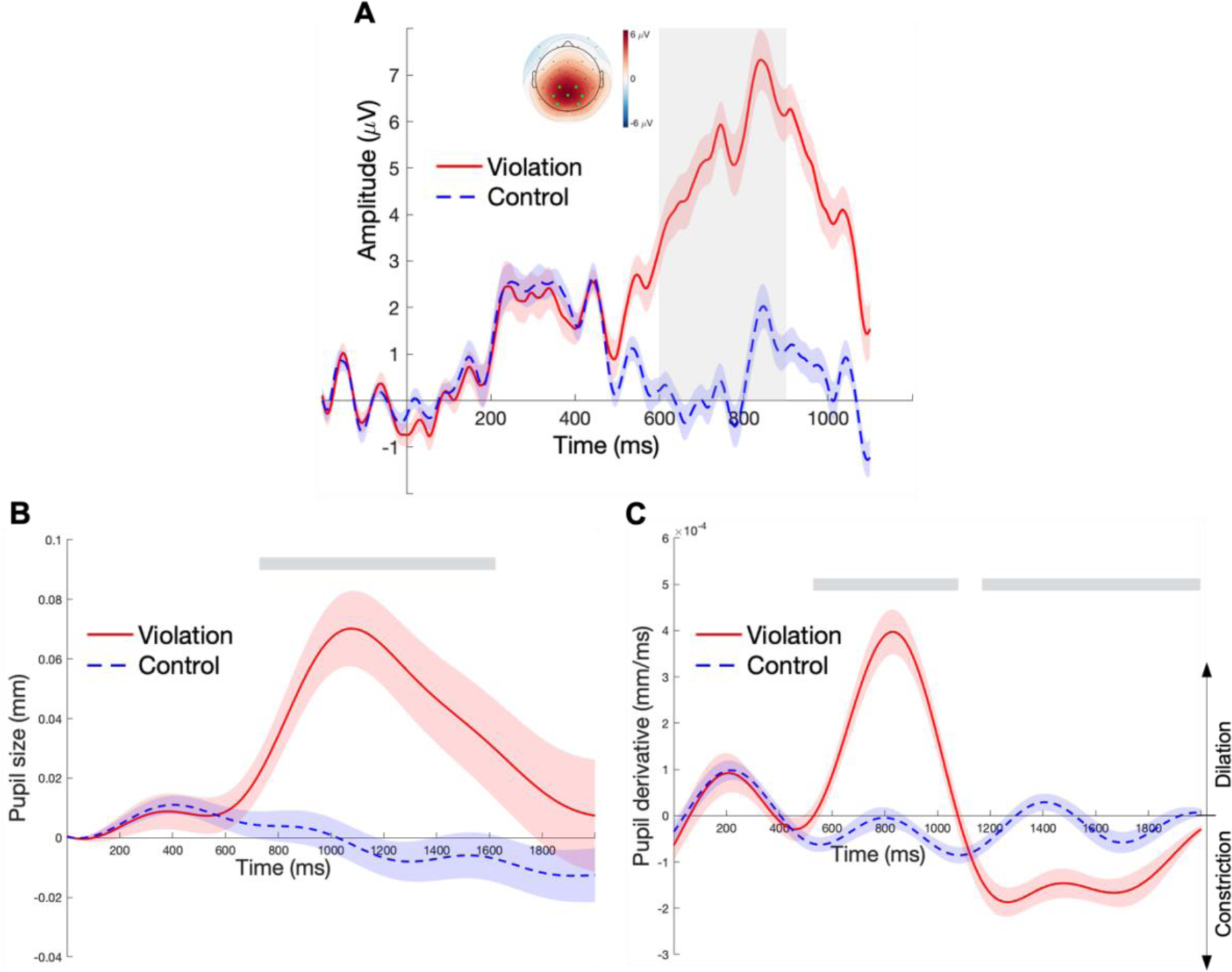
Grand averaged ERP amplitude (**A**), pupil size (**B**), and pupil derivative (**C**) for the two conditions of the sentence processing task, time-locked to the onset of the target word. **A**: Average amplitudes of the posterior ROI channel cluster (marked in light green on the topographic map). Trial-wise P600 amplitudes were calculated as the mean in the time interval marked by the grey box (600-900 ms). The topographic map shows the mean amplitude difference between violations and controls within the same time window. **B**: The grey bar highlights the time cluster with a significant difference in pupil size between the two conditions (730-1630 ms) and in which mean pupil size values were computed for each trial. **C**: The light grey bars highlight the time clusters for which the cluster-based permutation test revealed a significant difference between violations and controls in the pupil derivative (dilation at 540-1078 ms and constriction at 1170-2000 ms). The lefthand time window (dilation) was used to compute mean pupil derivative values for each trial in our statistical models. Error bands in A, B, and C indicate the group SEM based on subject-wise average time series.

#### 3.2.3 Relationship between ERP amplitude and TEPR

We further analyzed whether ERP amplitudes were positively predicted by the size of the pupillary response on a trial-by-trial basis. Regarding pupil size, the most conservative model including only violation trials was not statistically significant (*β* = 0.007, *SE* = 2.284, *t* = 0.286, *χ2* = 0.086, *p* = .769, RSI model, Fig. 5A). In the additional, preregistered analyses with trials from both conditions, the size of the pupil indeed significantly predicted ERP amplitudes (*β* = 0.071, *SE* = 0.018, *t* = 3.953, *χ2* = 13.903, *p* < .001). Hence, larger pupillary responses were generally accompanied by larger ERP amplitudes in the P600 time window (Fig. 5B). Note though, that this effect was not present in the additional exploratory model that included condition as a covariate (*β* = -0.008, *SE* = 0.014, *t* = -0.586, *χ2* = 0.332, *p* = .565). Additional, preregistered PCA analyses (see suppl. material) however also support the relationship between the P600 and pupil size within these time windows: P600 amplitude on a trial-by-trial basis was exclusively predicted by a component of the pupil peaking at around 1000ms. Figure 5C additionally shows the average correlation between the two measures over time and further indicates that the relationship is indeed strongest within the predefined time windows (dashed rectangle). Figure A5 in the supplementary material shows these cross-correlations over time for trials from each condition separately. They again demonstrate that trials from the violation, not control condition are driving the positive correlation.

**Figure 5.**
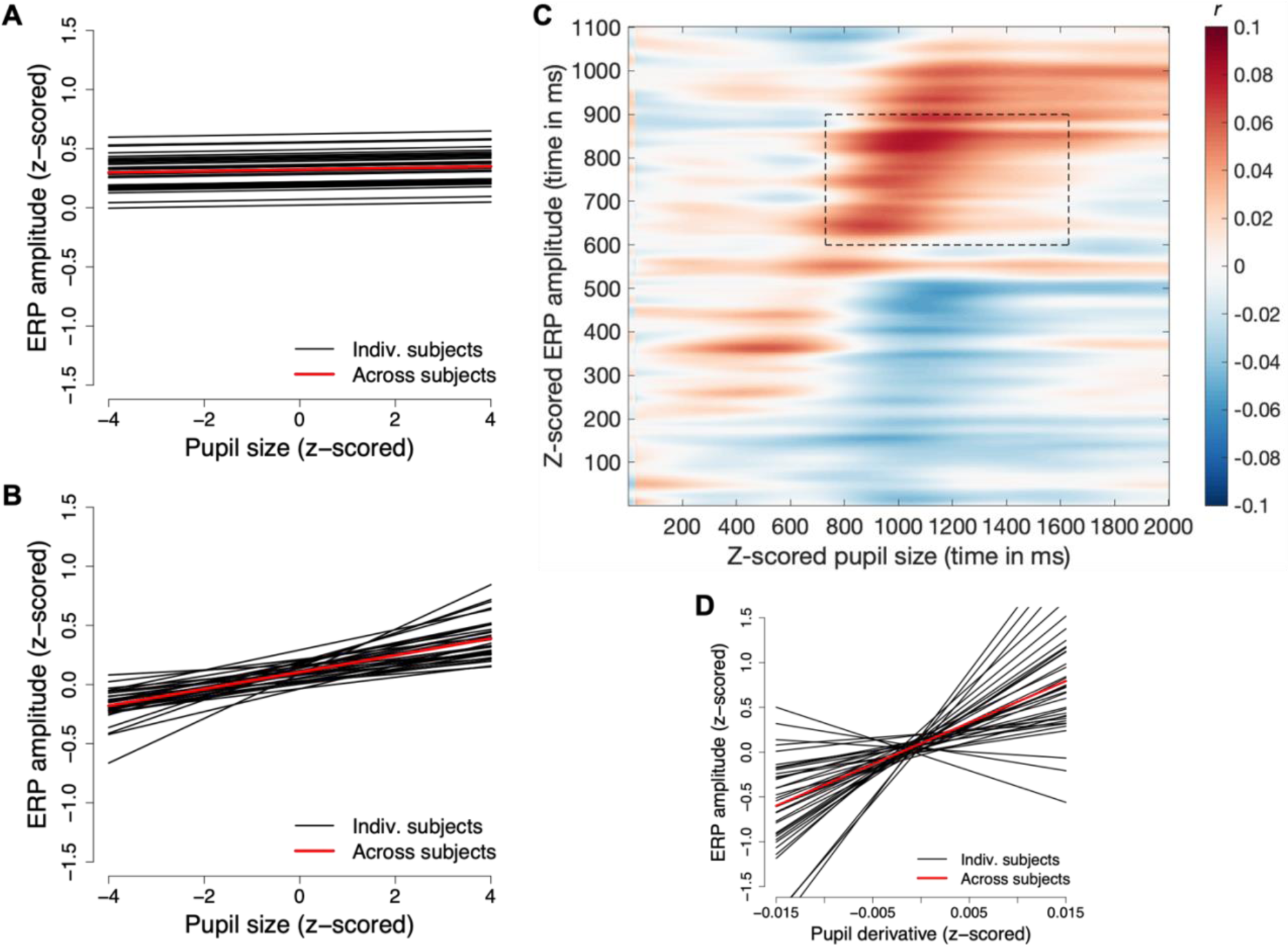
Relationship between the P600 and the TEPR in the sentence processing task. **A**: Predicted values for each participant and their mean from the main preregistered model testing the relationship between P600 amplitude and mean pupil size within the time window determined via the CBPT (light grey bar in Fig. 4B) within trials of the violation condition. **B**: Predicted values for each participant and their mean from the additional model testing the relationship between P600 amplitude and mean pupil size in the same, planned time window as in A, but within trials of both conditions. **C**: Cross-correlation of ERP amplitude and pupil size over time (i.e., average R of correlations within subjects). The dashed rectangle marks the combined time windows for analyses regarding the relationship between the two measures (i.e., for the model in A and B). **D**: Predicted values for each participant and their mean from the model testing the relationship between P600 amplitude and mean pupil derivative within trials of both conditions (in dilation time window indicated by the lefthand light grey bar in Fig. 4C).

Finally, additional, exploratory analyses revealed that the pupil derivative positively predicted ERP amplitudes on a trial-by-trial basis. Notably, this was not only found across all trials (*β* = 46.411, *SE* = 9.611, *t* = 4.829, *χ2* = 18.557, *p* < .001, see Fig 5D), but also for trials within the violation condition (*β* = 24.19, *SE* = 7.362, *t* = 2.039, *χ2* = 10.789, *p* = .001, RSI model).

#### 3.2.4 Relationship ERP-TEPR coupling between the two tasks

Finally, we explored whether a participant’s coupling between the pupil dilation and ERP component in one task was related to its coupling in the other task. We did so by correlating the respective participant coefficients of the ERP-pupil dilation models in the oddball and sentence processing task. Indeed, the size of the coupling across participants positively correlated across tasks, but again, specifically when considering the pupil dilation in the oddball task around the peak of the response where we also found the pupil-ERP coupling in the first place (*R* = .347, *p* = .038). Such a significant correlation was not detectable in the planned longer time window (*R* = 0.259; *p* = .127), where the ERP-pupil correlation was not significant in the first place (see section 3.1.3).

## 4 Discussion

In this study, we investigated whether the ERP components P3 and P600 might be linked to the phasic release of NE by relating them to the task-evoked pupillary response, a putative correlate of LC-NE activity (Breton-Provencher & Sur, 2019; Joshi et al., 2016; Liu et al., 2017; Reimer et al., 2016). We found that the pupillary response and the P3 component show a strikingly similar response pattern: Both the size of the pupillary response as well as ERP amplitudes were larger on rare oddballs than frequent standard stimuli (see also e.g., Friedman et al., 1973; Murphy et al., 2011; Qiyuan et al., 1985). For the first time, we also found these two measures to be related on a trial-by-trial basis when considering the pupil size around the peak of the response. In addition, our study is the first to demonstrate the same pattern for the P600 in a linguistic context: Just like the P600, the pupillary response was also larger on syntactic violations than correct control sentences and the ERP amplitudes could be predicted by pupil size on a trial-by-trial basis. Finally, we also found these exact patterns in both tasks regarding the positive derivative of the pupil, which suggests that it is specifically the dilation of the pupil that correlates with the P3 and P600. Our results thus suggest a close link between the neurophysiological and the pupillary response in *both* paradigms and regarding *both* late ERP positivities. Interestingly, the two effects (ERP-pupil coupling) even correlated across these two paradigms, suggesting a close relationship between the two ERP components and their link to NE. Combined, these findings further suggest that both components rely on a shared neural generator and, more specifically, that both may be linked to norepinephrine release in response to rare and motivationally significant stimuli (Nieuwenhuis, Aston-Jones, et al., 2005; Sassenhagen et al., 2014; Sassenhagen & Bornkessel-Schlesewsky, 2015).

Although the oddball effect on the pupillary response was replicated across studies, a relationship between the amplitude of the P3 and the size of the task-evoked pupillary response has not been found so far (Hong et al., 2014; Kamp & Donchin, 2015; LoTemplio et al., 2020; Murphy et al., 2011). Indeed, with our planned time window (based on the CBPT cluster with condition differences), the P3 and pupil size were not correlated either. It is important to note that this planned time window (as well as most time windows used in previous studies) was extensive, reaching until the very end of the epoch. However, pending replication, our further analyses actually indicated that the relationship regards a much more specific time window than previously assumed: Pupil size means around the peak of the dilation significantly predicted ERP amplitudes. This was also supported by both the early principal component of the pupil response which predicted P3 amplitudes and the early time period regarding the pupil size in which the two measures correlated in our cross-correlation plot. Crucially, analyses of the pupil derivative revealed that the actual dilation of the pupil for oddballs compared to standards indeed only happened much earlier and for a short period of time (within the first half of the epoch) and it is precisely this dilation that correlates with P3 amplitude.

Our results thus also suggest that there is a temporal disparity between the long-lasting oddball effect on pupil size and its early, transient correlation with P3 amplitude. It has been suggested that the pupil size following oddballs might be influenced by several factors, of which only one or some are related to the cognitive processing and others to the motor preparation, behavioral response, or light reflex (e.g., Steinhauer & Hakerem, 1992). Since the P3 oddball effect requires engagement in the task, all studies on the relationship between the pupillary response and P3 have used active paradigms, meaning that participants respond to stimuli and that these responses are systematically confounded with condition (either only one or a different button press for oddball targets). Thus, increased pupil size for oddballs compared to standards at the later stages of the epoch as observed here might have been caused by an additional effect of increased response preparation for the oddball or inhibition of the standard response (Hupé et al., 2009; Moresi et al., 2008). There was an effect of oddballs not just on the pupil but also on response (preparation) in that reaction times were longer for oddballs than standards. However, this response (preparation) might have not or even reversely affected the ERP amplitude (e.g., as influence of the negative-going readiness potential), diminishing any positive relationship between the pupil and P3 within the second half of the epoch, which makes it impossible to find a relationship with the P3 when considering the pupil size across the entire epoch. Future research might further elucidate this issue by using a delayed response or passive oddball paradigm.

How do the current findings further help us understand how the brain processes anomalies and ambiguities in the linguistic input and how this finally elicits the P600? The link between the P600 and the pupillary response that we found here adds further evidence to the proposal that the P600 typically observed in language paradigms might be part of the oddball-sensitive P3 family (e.g., Coulson et al., 1998; Kutas & Hillyard, 1983; Sassenhagen & Fiebach, 2019) and specifically reflects NE release from the LC (Bornkessel-Schlesewsky & Schlesewsky, 2019; Sassenhagen et al., 2014; Sassenhagen & Bornkessel-Schlesewsky, 2015). If this is indeed the case, we can learn much from the P3 about the P600 since the P3 is one of the most studied ERP components (see also Leckey & Federmeier, 2019). For example, we could transfer what we know about how it changes with age (e.g., Fabiani et al., 1998), how it can be used as a clinical marker (e.g., Polich & Herbst, 2000), and where it might be generated in the brain (e.g., Linden, 2005; Soltani & Knight, 2000). In addition, the P600-as-P3 view also revises theoretical considerations: Traditionally, the P600 has been interpreted as reflecting a language specific process where a sentence structure is built up or reprocessed, or the cost associated with these processes (e.g., Friederici, 2002; Hagoort, 2003; Osterhout et al., 1994). This view possibly evolved upon the initial observation of the P600 in response to syntactic violations and ambiguities (e.g., Friederici et al., 1993, 1996, 2001; Hagoort & Brown, 2000; Kaan & Swaab, 2003; Osterhout & Holcomb, 1992; Osterhout & Mobley, 1995). It has then become common to use the P600 as a marker of the presence or absence of such structural processing in different experimental designs. However, this reverse inference (Kappenman & Luck, 2011; Poldrack, 2006) might not be warranted. As outlined in the introduction, a plethora of studies has uncovered that the P600 is sensitive to all kinds of anomalies, even outside of language (e.g., Christiansen et al., 2012; Münte et al., 1998; Sitnikova et al., 2008). The present findings linking the P600 to the P3 further suggest a more parsimonious interpretation of the P600, in that it might reflect saliency processing more generally, possibly on different levels that are not always readily apparent. If the P600 is thus not as language-specific as previously assumed, this adds to the old, but ongoing debate on whether language processing recruits domain-general cognitive skills (e.g., Christiansen & Chater, 2008; Fedorenko, 2014; Fodor, 1983). In particular, a correspondence between the late positivities and the pupillary response suggests a crucial role for a neuromodulatory system that is highly general and not necessarily tied to a specific “domain” (Aston-Jones & Cohen, 2005; Sara, 2009). The diversity of anomalies that the P600 is sensitive to as well as its potential kinship to the P3 component also suggests that the underlying computational properties might be more domain-general. In fact, linking the P600 to neuromodulatory processes like NE could further help to understand how the brain handles salient and motivationally significant events in general, and how these processes are applied during listening or reading in particular. Specifically, it might promote alternative, explicit computational accounts of the P600 and its role in language processing, inspired by existing computational ideas on NE activity. For example, the process reflected by the P600 might be initiating model uncertainty as has been proposed for NE (Dayan & Yu, 2006; Muller et al., 2019). Further, the P600 might then be tightly linked to attention, both as a precondition to detect salient stimuli (Contier et al., 2022) as well as a consequence of NE projection.

Specifically, NE is suggested to lead to neural gain, that is, to increase the signal-to-noise ratio for processing of behaviorally relevant stimuli (Aston-Jones & Cohen, 2005; Nieuwenhuis, Aston-Jones, et al., 2005). Relatedly, attention might be re-oriented, as the NE possibly reflected in the P600 has been proposed as an “interrupt” signal that resets cortical networks (Bouret & Sara, 2005). In line with this idea, the P600 can be observed in response to disambiguating words that trigger a revision of the sentence interpretation (e.g., Kaan & Swaab, 2003; Osterhout & Holcomb, 1992).

In this study, participants actively judged whether sentences were ungrammatical or not, but the P600 is also elicited in linguistic tasks without such an overt task (Hagoort et al., 1993; Kaan et al., 2000). Is it possible that the P600 that is linked to the pupillary response is not “the” linguistic P600, but a separate, decision-related positivity (i.e., a P3 during language comprehension)? Although our data cannot entirely rule out this possibility and future research should test the link between the P600 and pupil dilation also using passive paradigms, there are several reasons why we think this is unlikely. First, if the observed positivity is indeed a P3, then the question arises where the P600 is. In other words, if the P600 observed here is actually a P3 that is linked to the pupil/NE, then this seems to be the relevant positivity elicited during language paradigms and is relevant to be investigated further. Secondly, a large portion of the P600 findings thus far is based on paradigms that employ sentence acceptability judgements (Allen et al., 2003; Gunter & Friederici, 1999; Hahne & Friederici, 1999; Kaan & Swaab, 2003; Kim & Osterhout, 2005; Osterhout & Mobley, 1995, just to name a few). Our finding of a link to the pupillary response thus pertains to the P600 as it has been studied in most of the previous literature.

If LC/NE activity is indeed involved in this linguistic saliency processing, then the question arises how the latency of the P600 is compatible with the fast responses of the LC/NE activity in response to salient stimuli (150-200 ms post-stimulus onset, Nieuwenhuis, Aston-Jones, et al., 2005). It is assumed that the LC/NE (and by extension, P3) response reflects the outcome of a decision-making process in which the stimulus is evaluated (Nieuwenhuis, Aston-Jones, et al., 2005) and its latency has been shown to vary with, for instance, familiarity of the stimulus categories (Aston-Jones et al., 1997). Only after the stimulus is evaluated to be salient and/or incongruent, the responsible cortices project to the locus coeruleus, which then initiates noradrenergic phasic responses, which again take up 100-150 ms to project to cortical target areas (Nieuwenhuis, Aston-Jones, et al., 2005). This is what is ultimately picked up by the EEG and indeed, the prototypical latency of the P3 thus fits into this idea. However, even the P3 often peaks much later, and both the P3 and P600 are very variable in latency, indeed depending on stimulus complexity (Kutas et al., 1977; Verleger, 1997) which presumably influences the decision-making process leading up to NE release from the LC/NE in the first place. In fact, despite being labelled as dichotomous “P3(00)” and “P600”, parietal positivities have been observed on a latency continuum. Only with very simple stimuli, such as flashing lights and auditory clicks as in the very first P3 study (Sutton et al., 1965), the positivity peaks as early as 300 ms. Many paradigms using slightly more complex stimulus categories such as symbols, heads, letters, or physical anomalies in written language (e.g., Kutas & Hillyard, 1980; Osterhout et al., 1996) elicit the positivity at around 400 ms. “Word” oddballs peak yet later, at around 500 ms (e.g., Kamp et al., 2015), which is in line with the general finding that orthographic processing and lexico-semantic access of a single incoming word already takes around 200 ms in the first place (Hauk et al., 2006). In sentences, combinatorial and semantic factors influence word processing 100-200 ms later (e.g., Kutas & Federmeier, 2011; Trueswell et al., 1993), explaining positivity latencies of 600-700 ms on violations and ambiguities such as here. The subsequent behavioral response, which positively correlates with the LC/NE phasic response (Bouret & Sara, 2004), has been shown to happen around the same time as the respective peak of the P3 in oddball and P600 in language paradigms (Sassenhagen & Bornkessel-Schlesewsky, 2015). In the same vein, we show here that also the pupillary response following violations in sentences is delayed compared to oddballs (by appr. 300-400 ms, around the same temporal divergence as between the peaks of the two ERP components). The time of both the behavioral and pupillary response thus mirrors this continuous “delay” in stimulus evaluation leading up to the release of NE. In fact, the temporal delay between the peak in the ERP and pupil dilation positively correlated across the two paradigms (*R* = .279, *p* = .044, see supplementary material for details on this additional, exploratory analysis). Thus, participants with a longer delay between the P3 peak and pupil dilation peak in the oddball paradigm also displayed a longer delay between the P600 peak and pupil dilation peak in the sentence processing paradigm. Future research could investigate these latency differences in the positivities varying with linguistic complexity more systematically, possibly combined with behavioral responses to link these temporal relationships more closely to the LC/NE system.

One question is how the observed relationship between the ERP components and the pupillary response relates to the oddball/violation effect. We found the trial-by trial correlation between ERP components (P3, P600) and the pupillary response in both the oddball and sentence processing task mainly when considering trial data from both the respective conditions together. In both tasks, both measures also showed the same effect of condition separately (i.e., larger response on oddballs and violations than standards and controls, henceforth generally called “oddball effect”). Should the relationship between ERPs and pupil be observed beyond this shared oddball effect? The hypothesis tested here suggests that both the ERP and pupillary response reflect a third, unobservable latent variable, namely LC/NE activity. The assumption is that the shared oddball effect actually “originates” in this LC/NE system, which is therefore only indirectly observable in both the ERP and pupillary response. Thus, the variance within the relationship between the two *should* mainly be driven by the shared oddball effect, which, in turn, should be equally observable in both measures, as is the case in this study. However, the issue remains that we cannot be completely certain whether the shared oddball effect is due to the link to the LC/NE system and not due to any other confounding variable. In a correlational study as the present one, we would need to show that variance beyond the binary, manipulated effect is also related between the two measures (i.e., correlated within separate conditions or when condition is controlled for). This is much more difficult within such a classic paradigm where stimuli were specifically designed to have large response variance between and low variance within conditions. This concerns especially variance presumably related to LC/NE activity within the critical oddball/violation condition, such as salience and probability. Given the simple task and stimuli, variance may even be lower in the oddball than sentence violation condition (which was indeed the case here when comparing within-participant standard deviations of ERP amplitude, *t*(35) = -14.389, *p* < .001). P3 oddball variance might be also reduced because it is thresholded in speeded tasks as the one here. In particular, it has been proposed that the P3 reflects the temporal built-up of evidence accumulation leading up to (and not the outcome of) decision making (e.g., Twomey et al., 2015). Such accumulation ends at a fixed threshold, which then triggers the response. This would naturally explain why shared response variance between the P3 and the pupil beyond the shared oddball effect is limited and why this is not such an issue in the sentence processing task, since the grammaticality judgement was not required until the end of the sentence. This lack of variance makes it even harder to identify any shared variance in our as well as previous oddball studies (e.g., Murphy et al., 2011).

Despite these difficulties, we still found evidence of such variance shared between the ERP amplitude and pupillary response beyond the oddball effect in *both* tasks: In the oddball paradigm, we could detect a relationship between P3 and peak pupil size even when controlling for the shared effect of task condition. Relatedly, the pupil PCA analysis of the oddball task revealed that the main, early principal component of the pupil predicted P3 amplitudes when focusing on oddball trials only. Similarly, in the sentence processing paradigm, a positive relationship between the pupil derivative and P600 amplitude was indeed detectable even when focusing on violation trials only. Lastly, ERP-pupil size cross-correlation plots from both tasks (see supplementary material) suggest that the positive relationship between the two measures in the critical time windows was mainly present in the oddball and violation condition. Taken together, both the shared oddball effect and the shared variance beyond it indicate that the P3 as well as P600 ERP components link to the pupillary response, and hence, possibly to the LC/NE system.

In the current study we used the pupillary response as an established index of LC/NE activity, however, as with many markers, the relationship between the pupil and NE involvement is still not completely understood. Several potential indirect connections exist between the two (Nieuwenhuis et al., 2011) and the pupil is most likely not solely influenced by norepinephrine, but also by other neuromodulators (e.g., serotonine, Cazettes et al., 2021; and acetylcholine, Larsen & Waters, 2018). Interestingly though, previous research suggests that while NE and acetylcholine both covary with pupil size, it is particularly NE that is also strongly correlated with the derivative of the pupil (Reimer et al., 2016), the measure that was strongly related to ERP components in both tasks in the present study, and showed a trial-by-trial correlation with P600 amplitudes even when just focusing on violation trials. In concert with previous evidence, especially the response time alignment of both the P3 and P600 (Sassenhagen & Bornkessel-Schlesewsky, 2015) as well as the LC/NE system (Bouret & Sara, 2004), our findings further suggest that the two components might be linked to NE. Future studies could corroborate these findings by linking the components to other NE correlates, for instance, salivary alpha amylase (e.g., Warren et al., 2017).

To circumvent the correlational problem due to the potential confound of condition as well as the issue with the indirect measure, the components’ link to NE would ideally also be complemented by evidence where NE is manipulated more directly, for example via stimulation or pharmacology (e.g., Joseph & Sitaram, 1989). Indeed, a non-invasive stimulation method, which acts upon the auricular branch of the vagus nerve projecting directly to the LC, has already been shown to enhance the amplitude of the P3 (Ventura-Bort et al., 2018), alpha amylase levels (Giraudier et al., 2022), and pupil dilation (D’Agostini et al., 2022; Sharon et al., 2021), revealing a promising approach to more directly test the involvement of the LC-NE-system in both ERP components (P3 and P600).

In conclusion, the present study suggests that the P3 and P600 are linked to the task-evoked pupillary response as a correlate of NE. Regarding the P3, our results specifically elucidate that the relationship to the pupil is temporally specific, potentially explaining mixed previous results on their relationship. We provide novel evidence that the late positivity in the linguistic context – the P600 – is also connected to the pupillary response, which supports the notion that the two ERP components might belong to the same family of positivities, thereby questioning the language-specificity of the P600. The link between the pupillary response and both late positivities suggests that these ERP components reflect a neuromodulatory process serving to increase or re-orient attention in response to salient and motivationally significant input.

## Conflict of interest statement

The authors declare no competing financial interests.

### Acknowledgements

We thank Marta Urbaniak, Lioba Bernd, and Antonia Heinrich for their help with stimulus preparation and data collection as well as Jan Ries for technical assistance. Funding was provided by the Research Focus Cognitive Sciences of the University of Potsdam (UFSKW) to Milena Rabovsky, Mathias Weymar, and Isabell Wartenburger.

## Footnotes

1 Note that regarding results, the following convention will signify the level at which analyses were planned: Without any specification, the respective analyses was included in the main hypothesis & analyses part of the preregistration. Analyses specified as “additional, preregistered” were mentioned in the exploratory part of the preregistration. Lastly, analyses specified as “additional, exploratory” were not part of the preregistration.

