## Supplementary material for "The P600 and P3 ERP components are linked to the task-evoked pupillary response as a correlate of norepinephrine activity"

### Oddball task

#### Principal component analysis of the TEPR (pupil size)

We preregistered that in case we would not find a statistically significant, positive relationship between ERP amplitude and the TEPR, we would decompose the pupillary response. To did so via single-trial principal component analysis using singular value decomposition and varimax rotation (thus technically, a factor analysis). Figure A1 displays the results of this analysis, namely the three components that combined explained more than 85% of the variance in the data.

**Figure A1.**

*Principal components of the pupil dilation in the oddball task. Percent of variance explained by each component in parentheses.*

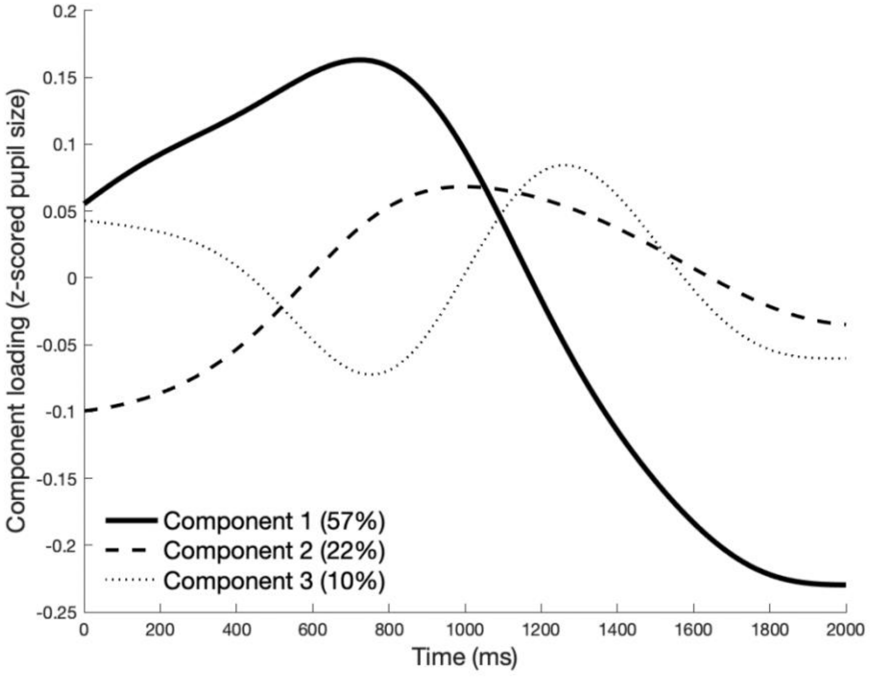

We then tested whether each component relates to the P3 amplitude, by testing whether each component's loading positively predicted ERP amplitudes on a trial-by-trial basis specifically within trials of the oddball condition. This was indeed the case for component 1 ($\beta$ = 2.693, *SE* = 0.816, *t* = 3.301, $\chi 2$ = 10.142, *p* = .001)[1]. In contrast, we failed to find such a significant effect on the P3 for either component 2 ($\beta$ = -1.778, *SE* = 1.004, *t* = -1.770, $\chi 2$ = 3.058, *p* = .08) or component 3 ($\beta$ = 1.610, *SE* = 0.835, *t* = 1.930, $\chi 2$ = 3.701, *p* = .054). Thus, within oddball trials, there is a positive relationship between the P3 amplitude and specifically component 1 of the pupil dilation, that is, the early part of the dilation, peaking at around 700 ms, but not later parts of the TEPR.

[1] Note that these results are based on linear mixed effects model predicting P3 amplitude by component loadings. To control for each component, we built one model including all three components as predictors simultaneously. Note that this model only converged with random slopes by participant for component 1, but not for component 2 or 3.

### Cross-correlation of ERP amplitude and pupil size: Oddball vs. standard condition

**Figure A2.**

*Cross-correlation of ERP amplitude and pupil size over time (i.e., average R of correlations within subjects) in the oddball task, separately for trials from the (**A**) oddball condition and (**B**) standard condition.*

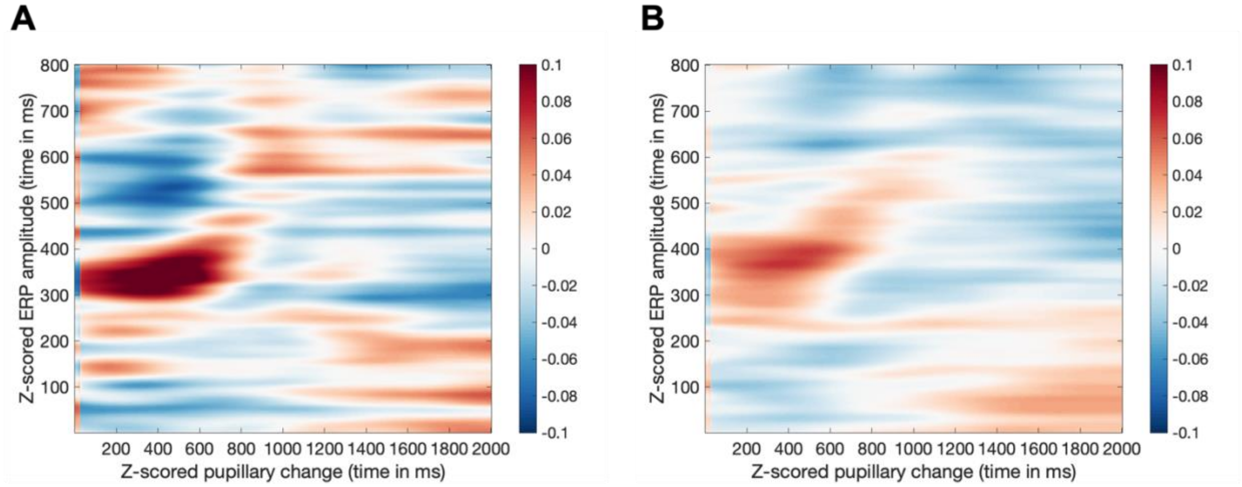


### Relationship between task-evoked signals and pupil baseline size

Phasic NE responses seem to exhibit an inverted U-shaped relationship to tonic (baseline) NE activity which is associated with arousal state (Aston-Jones & Cohen, 2005). In the medium tonic range, when one is attentive and engaged in the task, phasic bursts are large. However, when tonic mode is very low (drowsiness) or very high (distractiveness), phasic responses are small or absent. One possible index of tonic NE activity is baseline pupil size around the onset of the stimulus, and an oddball study by Murphy et al. (2011) indeed suggests that task-evoked pupillary responses show such an inverted U-shaped relationship to such pre-stimulus baseline pupil size. As pre-registered, we thus additionally tested whether the measures assumed to reflect phasic NE responses (P3 and TEPR) also show this inverted U-shaped relationship to baseline pupil size (0-50 ms relative to stimulus/target word onset).

Note that we (as e.g., Kuipers & Phillips, 2022) assumed that it is highly unlikely that we would get very small baseline pupil sizes in the first place, since participants are unlikely to be half asleep (which is also undesirable in EEG studies). Thus, we still considered a negative linear relationship between the ERP amplitude and baseline pupil size as indicating that the two might be related to phasic NE release (see also evidence by Hong et al., 2014).

However, concerns have been raised about relating the baseline-corrected TEPR to that baseline pupil size since these two measures are not independent of each other. For instance, regression to the mean (i.e., larger pupils tend to get smaller again and vice versa) can introduce an artifactual, negative relationship between the two which has little to do with their neurophysiological underpinnings (Mathôt & Vilotijević, 2022). These methodological issues impede any conclusions drawn from relating the TEPR to pupil baseline here (and subsequent comparisons between ERP components and pupil baseline) and we only report these results here for completeness because we pre-registered the corresponding analyses.

*Relationship between TEPR and pupil baseline size*

The model predicting the pupillary response (pupil size) by pupil baseline size indeed suggested a non-linear relationship between the two, considering pupil size in both time windows of interest (pre-registered time window based on cluster-based permutation test: $\beta_{lin}$ = -0.219, $SE_{lin}$ = 0.033, $t_{lin}$ = -6.660, $\beta_{quad}$ = 0.025, $SE_{quad}$ = 0.007, $t_{quad}$ = 3.697, $\chi 2$ = 13.665, $p$ < .001; exploratory time window around the peak: $\beta_{lin}$ = -0.132, $SE_{lin}$ = 0.034, $t_{lin}$ = -3.859, $\beta_{quad}$ = 0.016, $SE_{quad}$ = 0.006, $t_{quad}$ = 2.723, $\chi 2$ = 7.42, $p$ = .006)[2]. Note that $\chi 2$ and $p$ values are derived from the ratio likelihood test testing whether the non-linear model including baseline

[2] As pre-registered, these models as well as the following model testing the relationship between baseline pupil size and ERP amplitude included only data from the oddball condition. However, models including data from both conditions yielded the same results in all cases.

pupil squared explains the data better than the linear model. In both cases, the AIC was smaller for the non-linear model, additionally indicating a better fit (pre-registered time window: $AIC_{lin}$ = 3470.7, $AIC_{non-lin}$ = 3459; early, exploratory time window around the peak: $AIC_{lin}$ = 2855.2, $AIC_{non-lin}$ = 2849.8). However, the models revealed a rather U-shaped relationship between the two measures, hence pupil size was largest when baseline pupil size was either very small or very large (see Fig. A3A).

*Relationship between ERP amplitude and pupil baseline size*

The model predicting ERP amplitude (P3) by pupil baseline size suggested a positive, linear relationship between the two measures ($\beta$ = 0.015, *SE* = 0.006, *t* = 2.622, $\chi 2$ = 6.485, *p* = .011, AIC = 855.81, test against non-linear model: $\chi 2$ = 0.280, *p* = 0.597, $AIC_{nonlin}$ = 857.53; see Fig. A3B).

**Figure A3.**

*Predicted values of the models predicting (**A**) the TEPR (pupil size) and (**B**) P3 amplitude by baseline pupil size in the oddball task. Shaded areas display 95% confidence intervals.*

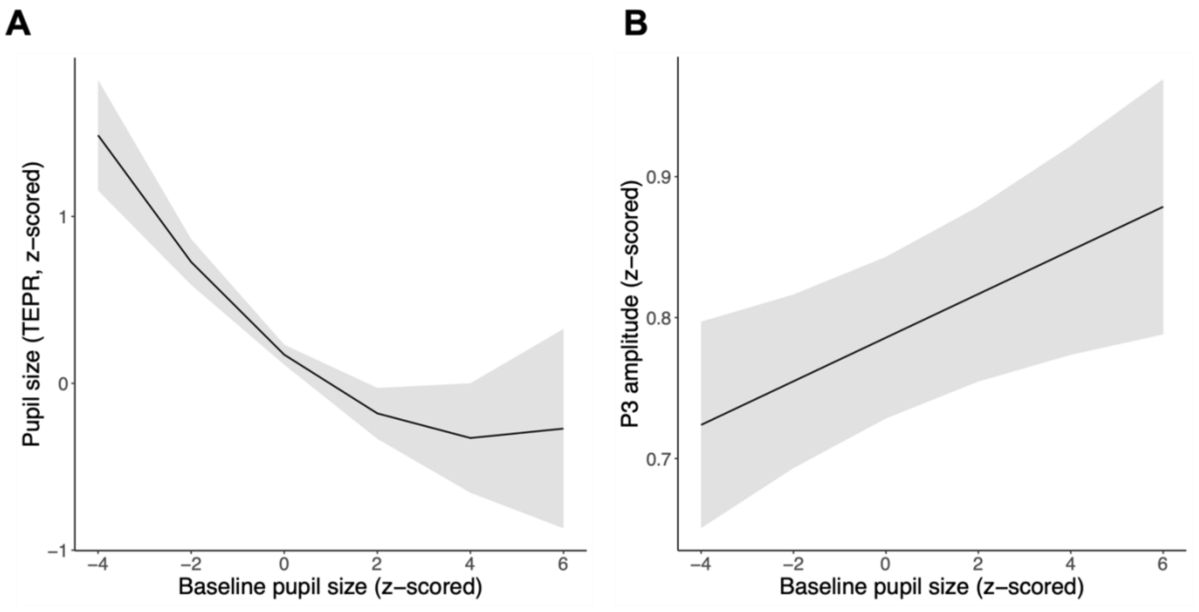

### Sentence processing task

#### Principal component analysis of the TEPR (pupil size)

As in the oddball task described above, we decomposed the pupillary response (pupil size) in the sentence processing task via single-trial principal component analysis using singular value decomposition and varimax rotation. Figure A4 displays the three components that combined explained more than 85% of the variance in the data.

**Figure A4.**

*Principal components of the pupil dilation in the sentence processing task. Percent of variance explained by each component in parentheses.*

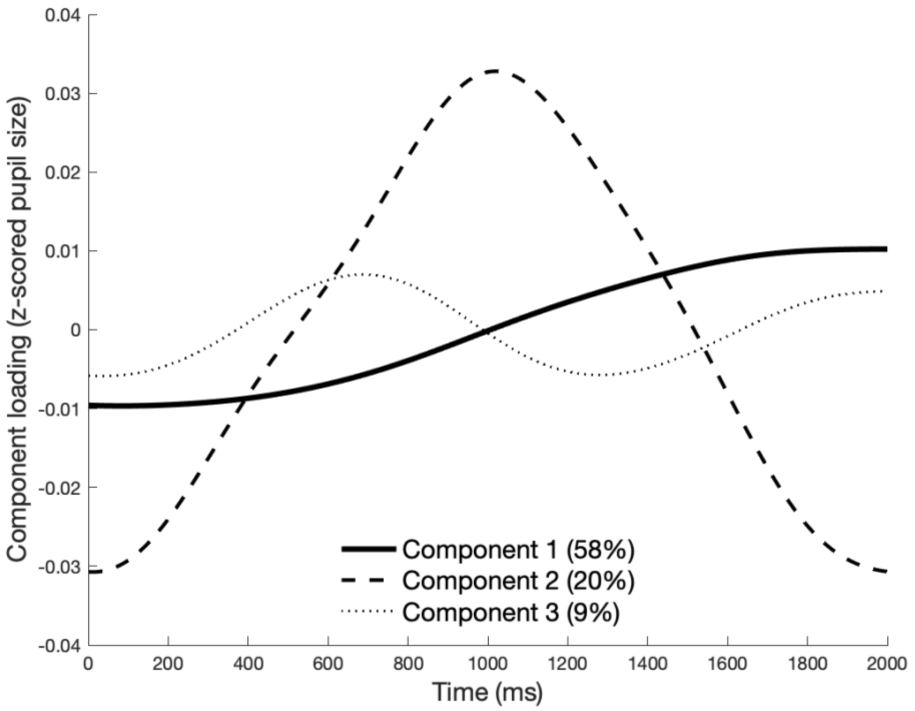


We again tested whether each component relates to the P600 amplitude, by testing whether each component's loading positively predicted ERP amplitudes on a trial-by-trial basis. The model including only violation trials failed to converge entirely, so we here report the results considering trials from both conditions together. Component 2 significantly and positively predicted P600 amplitudes on a trial-by-trial basis ($\beta$ = 2.568, *SE* = 0.704, *t* = 3.648, $\chi 2$ = 12.778, *p* < .001)[3]. In contrast, we failed to find such a significant effect on the P600 for either component 1 ($\beta$ = -0.553, *SE* = 0.548, *t* = -1.009, $\chi 2$ = 1.022, *p* = .312) or component 3 ($\beta$ = 0.119, *SE* = 0.533, *t* = 0.223, $\chi 2$ = 0.048, *p* = .827). This suggests a positive relationship between the P600 amplitude and specifically component 2 of the pupil dilation, that is, the part of the pupillary response that peaked around 1000 ms, which is in line with our results reported in the main text.

[3] Note that these results are based on linear mixed effects model predicting P600 amplitude by component loadings. To control for each component, we built one model including all three components as predictors simultaneously. Note that this model only converged with random slopes by participant for component 2, but not for component 1 or 3.

### Cross-correlation of ERP amplitude and pupil size: Violation vs. control condition

**Figure A5.**

*Cross-correlation of ERP amplitude and pupil size over time (i.e., average R of correlations within subjects) in the sentence processing task, separately for trials from the (**A**) violation condition and (**B**) control condition.*

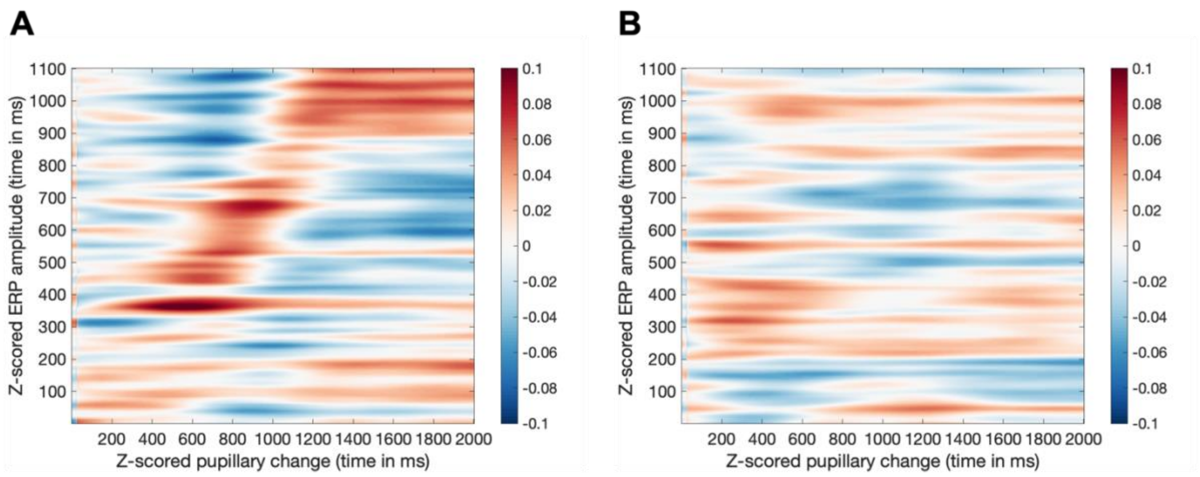


### Relationship between task-evoked signals and pupil baseline size

*Relationship between TEPR and pupil baseline size*

As in the oddball task, we hypothesized the pupillary response to exhibit an inverted U-shaped or negative relationship to baseline pupil size. The model predicting (task-evoked) pupil size by pupil baseline size in the sentence processing task suggested a non-linear relationship between the two ($\beta_{lin}$ = -0.151, $SE_{lin}$ = 0.023, $t_{lin}$ = -6.686, $\beta_{quad}$ = 0.046, $SE_{quad}$ = 0.007, $t_{quad}$ = 6.171, $\chi 2$ = 36.891, $p$ < .001)[4]. Note that $\chi 2$ and $p$ values are derived from the

[4] As pre-registered, these models as well as the following model testing the relationship between baseline pupil size and ERP amplitude included only data from the violation

ratio likelihood test testing whether the non-linear model incl. baseline pupil squared explains the data better than the linear model. The AIC value was smaller for the non-linear model, additionally indicating a better fit ($AIC_{lin}$ = 2598.1, $AIC_{non\text{-}lin}$ = 2563.2). Surprisingly, as indicated by the sign of the estimates, the model revealed a rather U-shaped relationship between the two measures, meaning pupil size was largest when baseline pupil size was either very small or very large (see Fig. A6A).

*Relationship between ERP amplitude and pupil baseline size*

The model predicting P600 ERP amplitude by pupil baseline size suggested neither a linear ($\beta$ = -0.003, *SE* = 0.012, *t* = -0.275, test against the null model: $\chi 2$ = 0.079, *p* = .779) nor a non-linear ($\beta$ = 0.006, *SE* = 0.008, *t* = 0.812, test against linear model: $\chi 2$ = 0.666, *p* = .415) relationship between the two (see Fig. A6B).

---

condition. However, models including data from both conditions yielded the same results in all cases.

**Figure A6.**

*Predicted values of the models predicting (**A**) pupil size (TEPR) and (**B**) P600 amplitude by baseline pupil size in the sentence processing task. Shaded area in A displays 95% confidence intervals.*

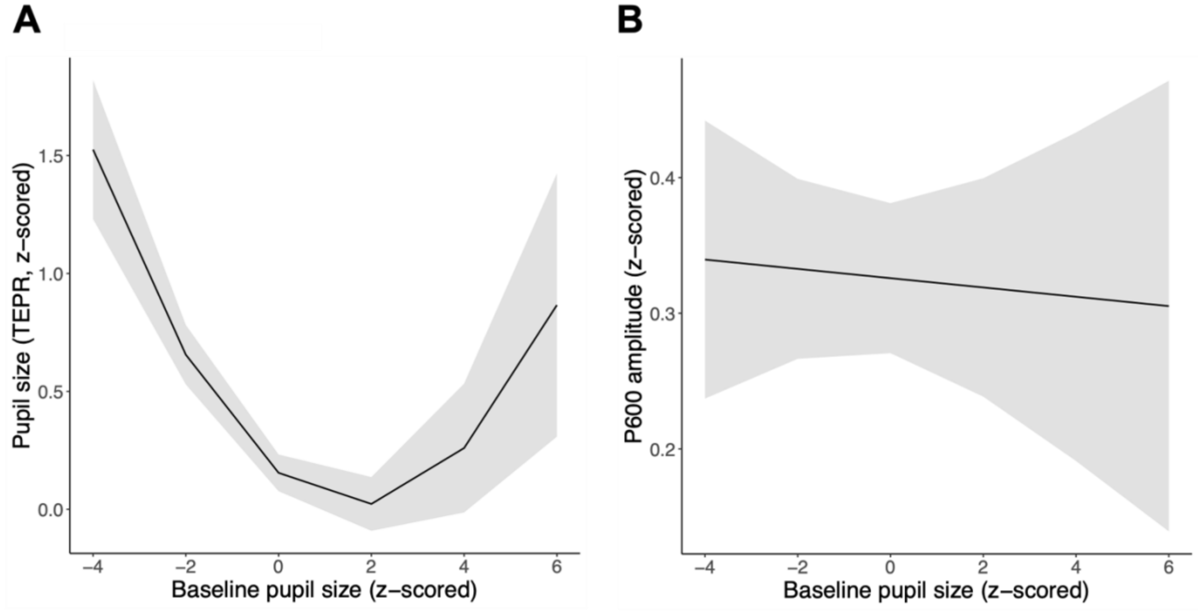

### Analysis details on correlation of temporal delay between ERP and pupil peak across paradigms

Latency of ERP component and pupil (size) peaks were obtained using the Jackknife resampling approach (e.g., Grenzebach et al., 2021; Kiesel et al., 2008; Luck, 2014; Sassenhagen & Bornkessel-Schlesewsky, 2015; Ulrich & Miller, 2001) on subject-wise condition difference wave time series. For each sampling of the Jackknife approach (grand average, each leaving out one subject average), we calculated the latency using 50% fractional area latency on positive grand average time series (i.e., negative values set to 0; Kiesel et al., 2008; Luck, 2014). These "peak" latencies were identified between 200-800 ms (P3), 200-1300 ms (oddball pupil), 400-1100 ms (P600) and 200-1600 ms (sentence processing pupil). Mean peak latencies in the oddball task were at 467.72 ms (*SD* = 2.037, P3) and 925.44 ms (*SD* = 3.121, pupil). Mean peak latencies in the sentence processing task were at 813.61 ms (*SD* = 1.961, P600) and 1157.2 ms (*SD* = 2.803, pupil). For each sampling, we then calculated the delay between the ERP peak and pupil peak in each task. Pearson correlation in concert with a permutation test (10 000 random permutations) indicated that these delays were positively correlated across the two tasks ($R = .279$, $p = .044$).
